# Identification of Small Molecule Dimethyoxyphenyl Piperazine Inhibitors of Alpha-Synuclein Fibril Growth

**DOI:** 10.1101/2025.06.26.661821

**Authors:** Helen Hwang, Dhruva D. Dhavale, Sarah J. Wang, John M. Beale, Nigel J. Cairns, Paul T. Kotzbauer

**Author notes:** Corresponding author: Helen Hwang, Department of Neurology, Washington University School of Medicine, Campus Box 8111, 660 South Euclid Avenue, St. Louis, Missouri 63110, USA.

## Abstract

Identification of Small Molecule Dimethyoxyphenol Piperazine Inhibitors of Alpha-Synuclein Fibril Growth Alpha-synuclein (asyn) fibril accumulation is the defining feature of Parkinson disease and is a target for disease-modifying treatments. One therapeutic strategy to reduce fibril accumulation is inhibition of asyn fibril growth. We developed a sensitive fluorescence-based fibril growth assay to screen for small molecule inhibitors. After validating the inhibition assay using a previously identified inhibitor, epigallocatechin-3-gallate, we identified compound 1 as a lead for inhibition of fibril growth. We analysed structure-activity relationships with analogs of 1 to optimize inhibition potency. Our results identified two dimethoxyphenyl piperazine analogs with more potent inhibition of in-vitro assembled fibrils. These analogs also inhibited the growth of asyn fibrils amplified from Lewy Body Disease brain tissue, further validating the inhibitor screening assay. Molecular docking studies indicate that these compounds can bind to the fibril ends, suggesting a potential capping mechanism through which these compounds inhibit the sequential association of monomeric asyn required for fibril growth.

## Introduction

Alpha-synuclein (asyn) is a small 140 amino acid protein that regulates synaptic vesicle-mediated protein trafficking^1^. In its native form, asyn is a disordered soluble monomeric protein. The N-terminus forms an amphipathic alpha-helix upon interactions with membranes. In the formation of fibrils, asyn adopts a fibrillar cross-beta sheet structure, which is considered pathologic^2–4^. Misfolding of asyn into beta-sheet amyloid fibrils coupled with accumulation is linked to the pathophysiology of Parkinson’s disease (PD) and Lewy body dementia (LBD), the dementia associated with PD^5^. Furthermore, dominantly inherited mutations in the gene encoding asyn (SNCA) have been identified in rare forms of hereditary parkinsonism that are also characterized by asyn fibril accumulation, supporting asyn fibril accumulation as a therapeutic target^6–8^.

Asyn fibril accumulation involves an initial nucleation event, where two or more asyn monomers come together to form a misfolded oligomeric seed or protofibril. Nucleation is followed by the growth phase or elongation phase, where the sequential addition of asyn monomers leads to the formation of fibrils with beta-sheet structure. Previous studies have utilized several approaches to identify inhibitors for asyn fibril nucleation and growth^9–16^. Stabilizing the monomeric form of asyn by DNA aptamers has been shown to delay fibril formation^17^. Others have shown that some polyphenols, including (−)-epigallocatechin gallate (EGCG) can redirect monomeric a-synuclein to form off-pathway spherical oligomers and thus inhibit beta-sheet fibril formation^18–26^. Dopamine and other catecholamines can similarly promote the formation of non-fibrillar oligomeric forms of asyn^27–30^. Other molecules, such as select antibiotics^28,31^, pigments^30,32–34^, glucosides^35–37^, quinones^38,39^, and pirimido-pyrazine^40,41^ also have the potential to inhibit asyn aggregation.

Conventional in vitro screening approaches to identify inhibitors of Asyn fibril accumulation commonly utilize a non-seeded aggregation approach^9,11,14,30^. One study adopted a seed amplification approach with repeated incubation and sonication to achieve exponential growth^42^. Both types of studies typically use high monomeric asyn concentrations (25-140 μM), continuous orbital rotation, and lengthy incubation time (16-48 hours) ^9,42^. We hypothesize that isolating and targeting the fibril growth pathway may yield additional candidate inhibitors for drug discovery.

Asyn fibril accumulation is typically quantified by measuring Thioflavin T (ThioT) fluorescence, either continuously or at an endpoint after a 24-36 hour incubation period. ThioT is a fluorescent dye that is weakly fluorescent in the presence of monomeric asyn but undergoes a substantial increase in fluorescence intensity by several orders of magnitude upon binding to amyloid fibrils, including asyn fibrils. We previously developed a fluorescence-based fibril growth assay that uses fluorescein arsenical hairpin binder (FlAsH) to detect the association of two or more asyn monomers with bicysteine tags^43,44^. The FlAsH assay relies on quiescent incubation of monomeric asyn with a bicysteine (C2) tag and wildtype asyn fibril seeds with no C2 tag. Furthermore, the incubation time is short (3 hours), during which the rate of fibril growth is constant. Since wild-type (WT) asyn fibril seeds produce no FLASH fluorescence, the fluorescence signal is specific to fibril elongation with C2-asyn. We previously observed similar growth kinetics with C2-asyn monomer and WT-asyn monomer, using ThioT as a readout, suggesting that addition of a bicysteine tag on the N terminus of asyn does not interfere with fibril growth. Given the higher sensitivity of FlAsH dye compared to ThioT fluorescence, this approach detects fibril growth over shorter periods of time (3 hours) compared to ThioT^43^.

We further developed the FlAsH fibril growth assay to screen small molecule compounds by using lower concentrations of monomer and fibril seeds and a short incubation period. We validated that this fibril growth assay measures an inhibitory effect of EGCG on fibril growth as observed in previous studies with ThioT and then used this fibril growth assay to identify lead compound **1**. To identify more potent inhibitors, we performed a series of structure-activity testing and identified two analogs of **1** that were more potent inhibitors. These analogs of **1** also inhibited the growth of asyn fibrils amplified from postmortem Lewy Body disease (LBD) brain tissue.

## Results

### An In Vitro Asyn Fibril Growth Assay for Screening Small Molecule Inhibitors

Inhibition of fibril growth by small molecule compounds depends on binding interactions with either monomer, fibrils, or both. In one proposed mechanism, a compound binds to the end surface of fibrils and interferes with the binding of the new monomer required to extend the seed (Figure 1). Since binding will deplete the free concentration of the inhibitor, measured IC_50_ values will be proportional to the concentration of the binding target (fibril seeds) and proportional to the concentration of monomer, with which the inhibitor competes for binding. Thus, we sought to optimize the FlAsH fibril growth assay by decreasing the concentrations of monomer and fibril seeds and found that a combination of 1 μM fibril seeds, 3 μM monomer, and 3-hour incubation time provided sufficient signal to analyze dose-response curves and determine IC_50_ values.

**Figure 1.**
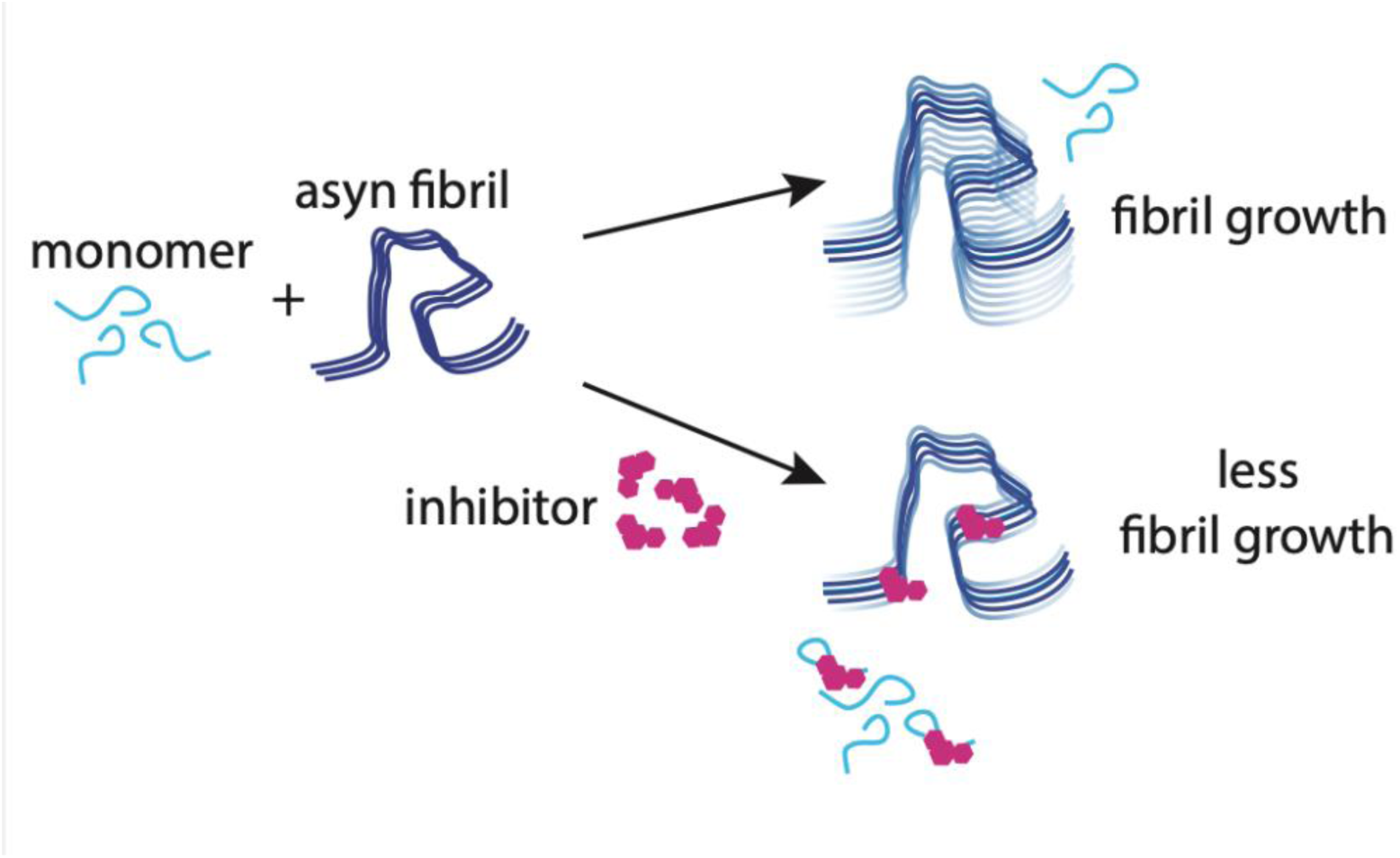
Schematic of fibril growth inhibition. Proposed mechanism of inhibition of asyn fibril growth by a small molecule inhibitor (pink) binding to the end surface of fibrils.

We utilized the optimized FlAsH assay to evaluate a previously identified fibril growth inhibitor, EGCG, and observed dose-dependent inhibition with an inhibition constant (IC_50_) of 8 ± 1 μM (Figure 2A). Interestingly, control reactions containing EGCG with no seeds showed increased fluorescence signal at high EGCG concentrations. This may correspond to the formation of off-pathway oligomers previously observed when asyn is incubated with EGCG^18,23^. We then tested a compound we previously identified as a potential ligand for asyn in molecular docking studies, ethyl 1-[(4-hydroxy-3,5-dimethoxyphenyl)methyl]piperidine-4-carboxylate (**1**), and observed that **1** inhibits fibril growth with an IC_50_ of 18 μM (Figure 2B and Figure 2C). In contrast to EGCG, the fluorescence signal did not increase in the control reactions of **1** with no added fibril seeds (monomer only), indicating the absence of oligomer formation or spontaneous nucleation of fibrils over the 3-hour incubation period.

**Figure 2.**
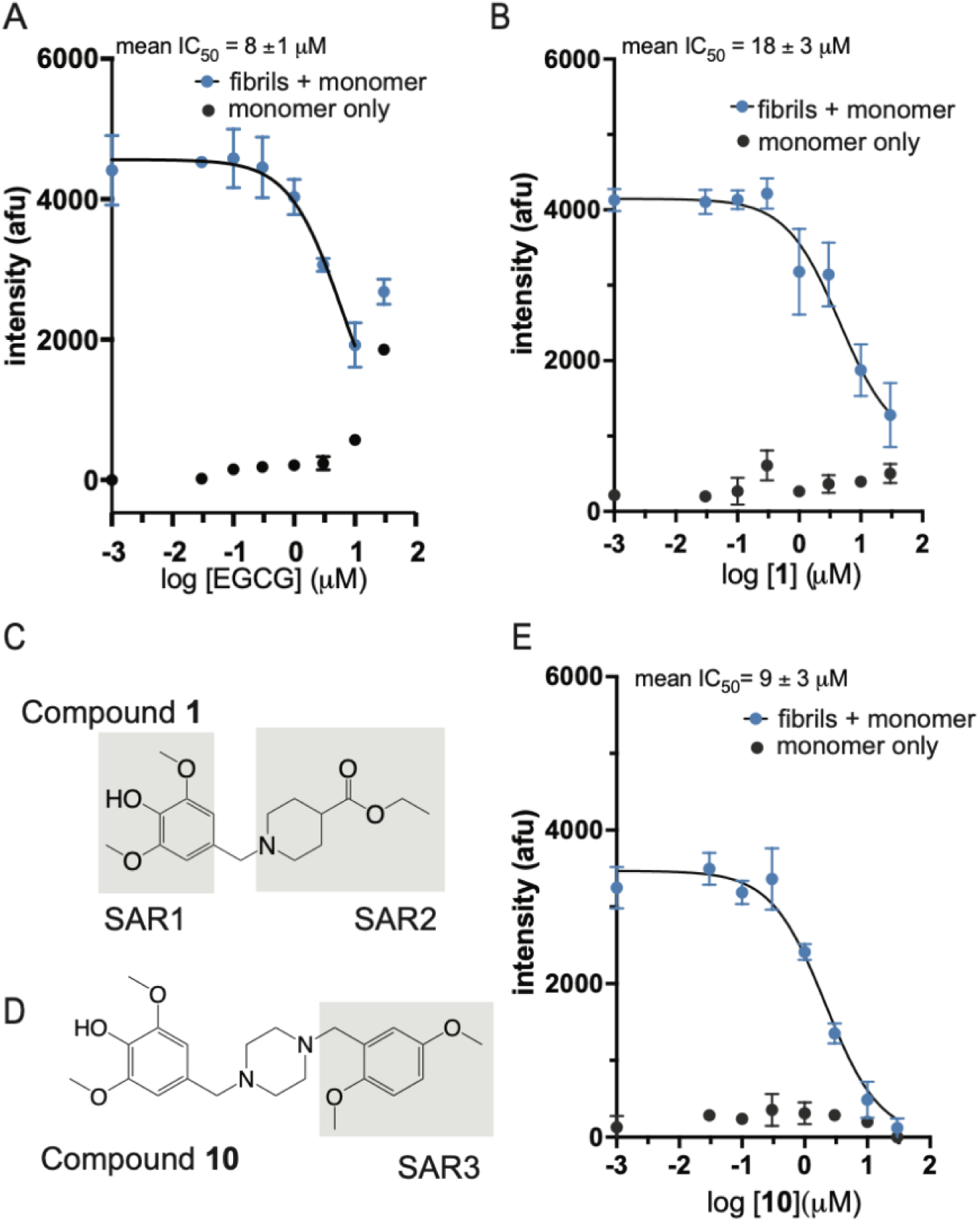
Inhibition of fibril growth by EGCG and newly identified compounds. A) Representative dose-response curve for EGCG in an optimized fibril growth assay based on FlAsH fluorescence. Note the increased fluorescence signal for monomer plus EGCG at 30 μM concentration, limiting the measurement of fibrils at this concentration. B) Representative dose-response for **1** in the optimized fibril growth assay. The fluorescence signal is low in the presence of monomer only and does not increase significantly with increasing compound concentrations. C) Chemical structure of **1** with shaded areas indicating regions where modifications were tested for structure-activity relationship (SAR) studies. D) Chemical structure of **10** with shading that indicates the region that was modified for the third set of structure-activity relationship studies. E) Representative dose-response for **10** in the optimized fibril growth assay. The fluorescence signal is low in the presence of monomer only and does not increase significantly with increasing compound concentrations. Data points represent mean ± standard deviation (std) (triplicates; n = 3 independent experiments).

### Screening additional analogs identifies structure-activity relationships for fibril growth inhibitors

To determine whether we can further improve the inhibition potency of compound **1**, we examined 29 analogs of **1** in three rounds of structure-activity relationship (SAR) analysis. First, we examined several analogs (**2-4**) with alternate substitutions in the benzyl group on the left-hand side of **1** (Supplementary Figure 1A, SAR1). We observed a loss of inhibition potency for all of these analogs. Next, when the piperidine in **1** was replaced with a piperazine and the ester was replaced with a chlorobenzyl-piperazinyl (**5**), the inhibition improved to 15 μM, and max inhibition was 99% compared to 18 μM and 79%, respectively, for **1**. Other substitutions of **1** with another piperazinyl or piperidine (**6 and 7**) did not improve fibril growth inhibition (right side, SAR2, Supplementary Figure 1B). We subsequently screened 23 piperazine analogs (**8-30**) with modification on the right-hand side and found several dimethoxyphenol piperazine compounds (**8-13**) that were more potent (lower IC_50_) and had greater maximum inhibition than **1** (Table 1). For example, **10**, a piperazinyl dimethoxylphenol ethanedioate, had an IC_50_ of 9 ± 3 μM (Figure 2D and Figure 2E). The analogs of **1** can be further divided into classes of phenacyl (**8**), pyridinyl (**9**), dimethoxy benzyl (**10**, Figure 2D), methoxy benzyls (**11** and **13**), and halogen benzyls (**5** and **12**). Substitutions that included bulkier subgroups (**14, 15, 18, 19, 20, and 21**) and thiol-containing analogs (**28, 29**, and **30**) had little inhibitory effect (Supplementary Figure 2). We also confirmed that 100 μM solutions of the test compounds had less than 15% higher native fluorescence relative to buffer alone, at the FlAsH excitation/emission wavelengths (Supplementary Table 1). We further evaluated whether apparent inhibitory effects were due to the fluorescence quenching of the FlAsH dye by the test compounds (Supplementary Table 2). Only two of the compounds (**6** and **15**) had quenching of more than 25% fluorescence in the presence of fibrils pre-assembled in vitro with C2-asyn monomers and incubated with FlAsH, but these two compounds showed minimal inhibition of fibril growth (Supplementary Figure 1B and Supplementary Figure 2). This further confirmed that the inhibitory effects observed for the compounds were not due to quenching of FlAsH dye by the compounds at higher concentrations.

**Table 1.**
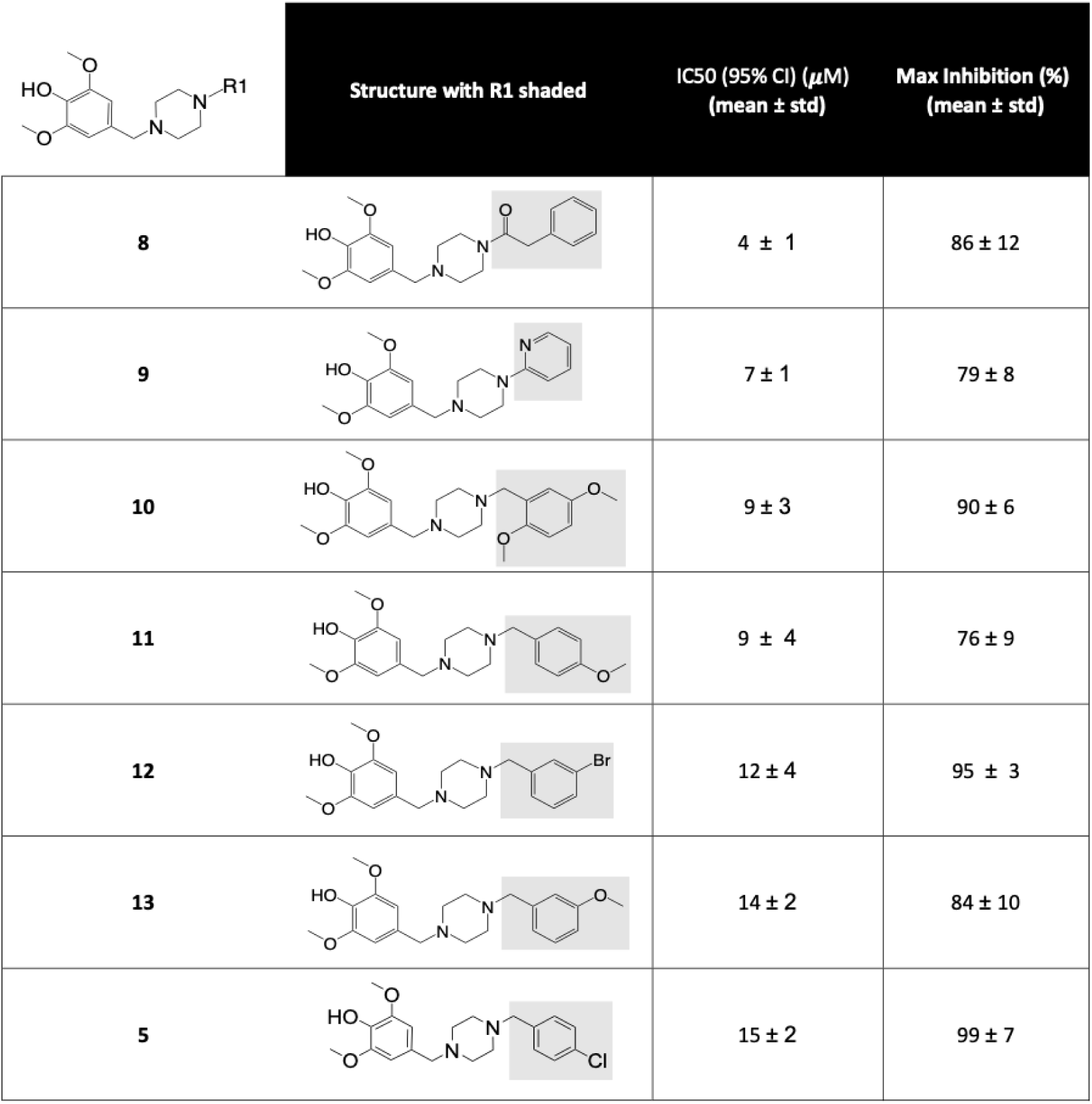
Compounds with highest inhibition potency after 3 rounds of structure-activity screening. . The structures share a piperazinyl and a 4-hydroxy-3,5-dimethoxybenzylgroups. IC_50_ is the mean ± standard deviation (std) and maximum inhibition is calculated as a (Intensity_100μM_-Intensity_0μM_)/intensity_0μM_ from three independent experiments.

|  |  | Structure with R1 shaded | IC<sub>50</sub> (95% CI) (μM)<br>(mean ± std) | Max Inhibition (%)<br>(mean ± std) |
| --- | --- | --- | --- | --- |
| 8 |  |  | 4 ± 1 | 86 ± 12 |
| 9 |  |  | 7 ± 1 | 79 ± 8 |
| 10 |  |  | 9 ± 3 | 90 ± 6 |
| 11 |  |  | 9 ± 4 | 76 ± 9 |
| 12 |  |  | 12 ± 4 | 95 ± 3 |
| 13 |  |  | 14 ± 2 | 84 ± 10 |
| 5 |  |  | 15 ± 2 | 99 ± 7 |

### Fibril Growth Inhibition Confirmed by Conformation-Specific Immunoassay

To confirm the inhibition of fibril growth for compounds **8** and **10**, we used an orthogonal fibril growth assay that does not rely on fluorescence. We developed a sandwich enzyme-linked immunosorbent assay (ELISA) that utilizes a conformation-specific antibody (Abcam 209538, MJFR-14-6-4-2) ^45^ in the capture step and an in-house monoclonal antibody 13G5 ^46^ to measure fibril concentration at the end of the growth assays. This immunoassay is approximately 300-fold more sensitive for detecting fibrils than monomers (Supplementary Figure 3). In contrast to the FlAsH assay, this immunoassay measures the level of fibril seeds at baseline, and growth is measured as an increase in signal relative to the baseline signal of fibril seeds. We increased the incubation time for the fibril growth reaction to 20 hours and increased the asyn monomer concentration to 6 μM, in order to increase the amount of fibril growth and compensate for the reduced sensitivity of the immunoassay relative to the FlAsH assay. Under these conditions, fibril concentrations increase by 70% to 100% over 20 hours in the absence of inhibitor compounds. We confirmed that the presence of the compound decreased fibril growth as measured by the ELISA in a concentration-dependent manner for compounds **8 and 10** (Figure 3A). FlAsH fluorescence measurements in parallel reactions showed similar inhibition patterns. (Figure 3B). Thus, the orthogonal assay provides confirmation that compounds **8** and **10** inhibit fibril growth and that FlAsH can successfully identify inhibitory compounds.

**Figure 3.**
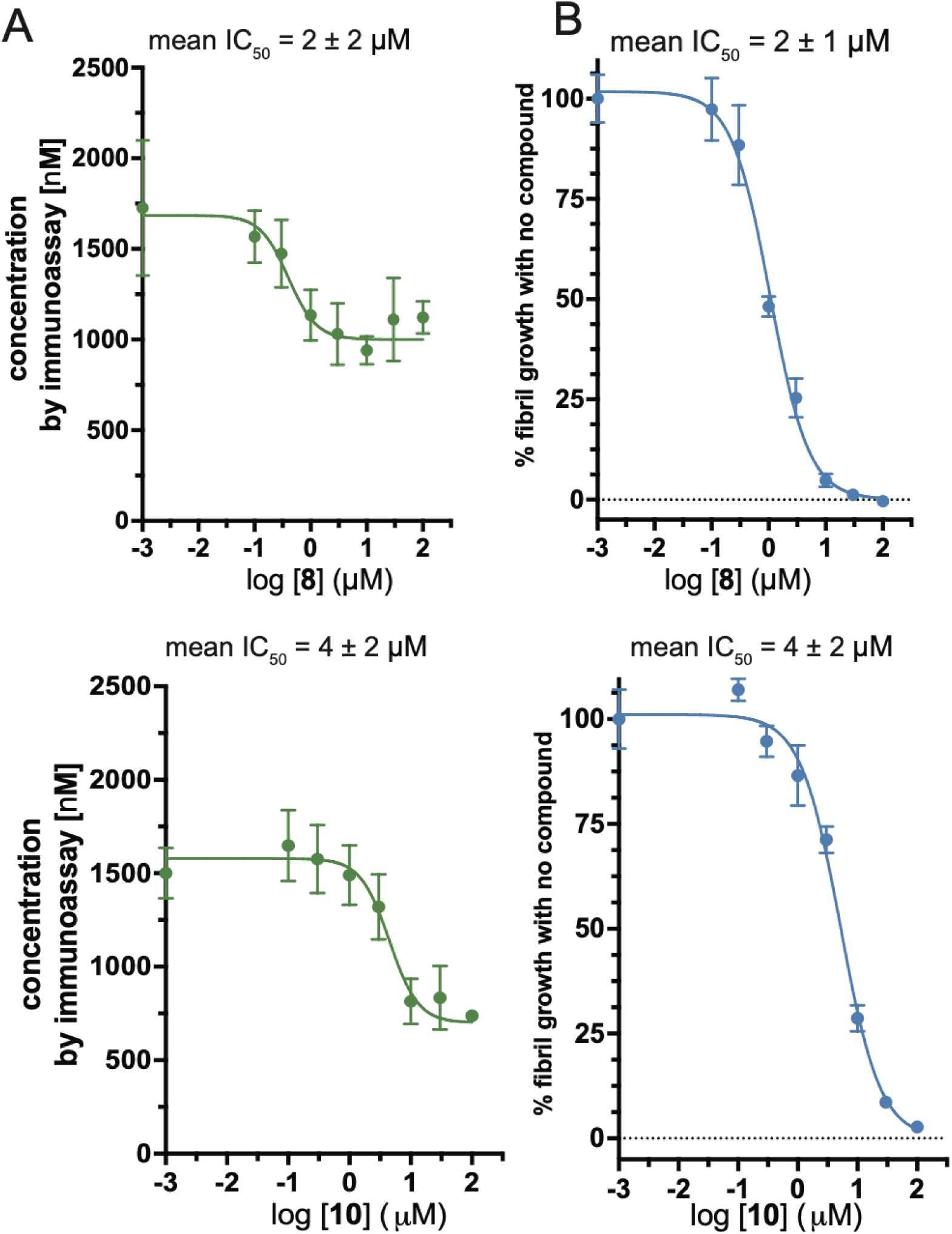
An immunoassay specific for asyn fibrils confirms that compounds inhibit fibril growth. A) Dose-response curves for **8** (top) and **10** (bottom) using a conformational-specific immunoassay to measure inhibition of fibril growth. B) Corresponding dose-response curves for **8** (top) and **10** (bottom) using a FlAsH fluorescence assay to measure inhibition of fibril growth; IC_50_ is mean ± std from two independent experiments. Note that the immunoassay detects both seeds and new fibril growth while the FlAsH assay detects only new fibril growth, accounting for differences in the apparent maximum reduction in signal by compounds.

### Fibril Growth Inhibition Observed by ThioT Fluorescence

We further evaluated whether fibril growth inhibition by our lead compounds can be measured by ThioT fluorescence, which has been widely used in previous studies of fibril growth inhibitors. Due to the low sensitivity of ThioT, we increased our fibril seed concentration to 3 μM and utilized continuous shaking for 20 hours to achieve measurable fibril growth. In our initial control experiments, we observed that compounds **8** and **10** reduced ThioT fluorescence in the presence of fibrils (Supplementary Figure 4A), which may be due to either quenching or competition with ThioT binding to fibrils. To address the potential for this effect to interfere with measurements of fibril growth, we used centrifugation and washing to isolate fibrils from compounds following the growth reaction. We then resuspended the fibril pellet in buffer containing ThioT (18 μM) to measure fibril growth. This protocol substantially reduced the quenching or inhibitory effect of the compounds on ThioT fluorescence (Supplementary Figure 4B), and we observed that compounds **8** and **10** significantly inhibited fibril growth (Figure 4).

**Figure 4.**
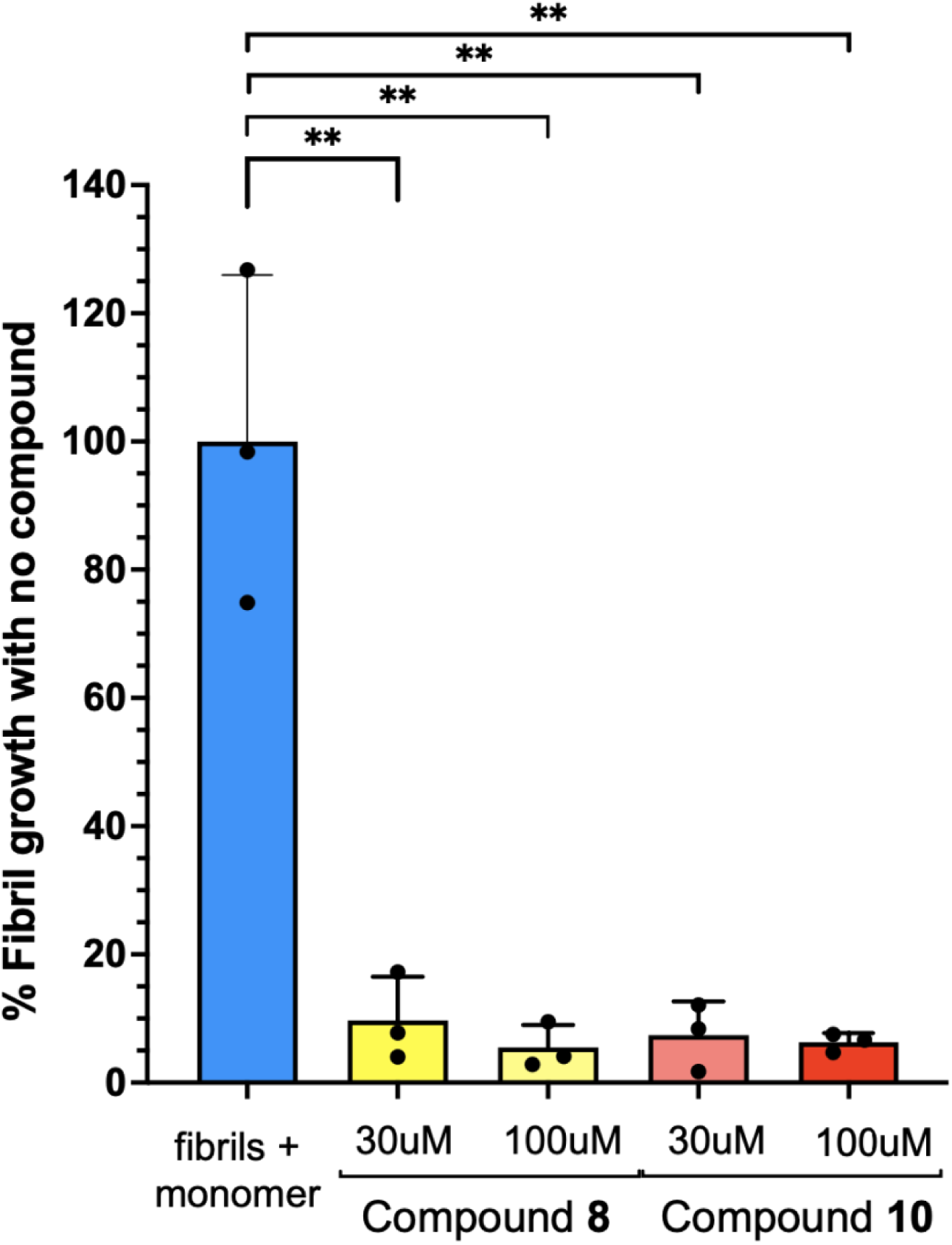
ThioT fluorescence shows inhibition of fibril growth by compounds. Measurement of fibril growth by ThioT fluorescence in the presence of **8** and **10 at** 30μM and 100μM respectively, after serial centrifugation and washing. There is significant fibril growth inhibition seen at both concentrations of **8** and **10** (t-test, p<0.01). Graphs show data points (n=3) with mean (bars) and std (error bars). Similar results were observed in three independent experiments.

### Dimethoxyphenyl Piperazine Compounds Also Inhibit the Growth of Fibrils Amplified from LBD Postmortem Brain Tissue

Emerging studies have shown that the structure of asyn fibrils derived from postmortem LBD brain tissue differs substantially from the structures of fibrils produced in vitro from recombinant protein^44,47^. Differences in structure among these distinct fibril conformers may affect the binding and inhibition of fibril growth by small molecule compounds. Compounds **8 and 10** were initially identified based on in vitro fibrils. We recently generated amplified LBD fibrils using fibrils extracted from LBD brain tissue, and showed that the fold of the LBD amplified fibrils matched the fold of asyn fibrils directly extracted from PD and LBD brain tissue^48^. We utilized amplified LBD fibril seeds in the same fibril growth assay to test these lead analogs^44^. Compounds **8** and **10** showed similar IC_50_ values to those obtained for in-vitro assembled fibril seeds (Figure 5). This result indicates that compounds **8** and **10** can inhibit the growth of fibrils that are structurally similar to those accumulating in PD and LBD.

**Figure 5.**
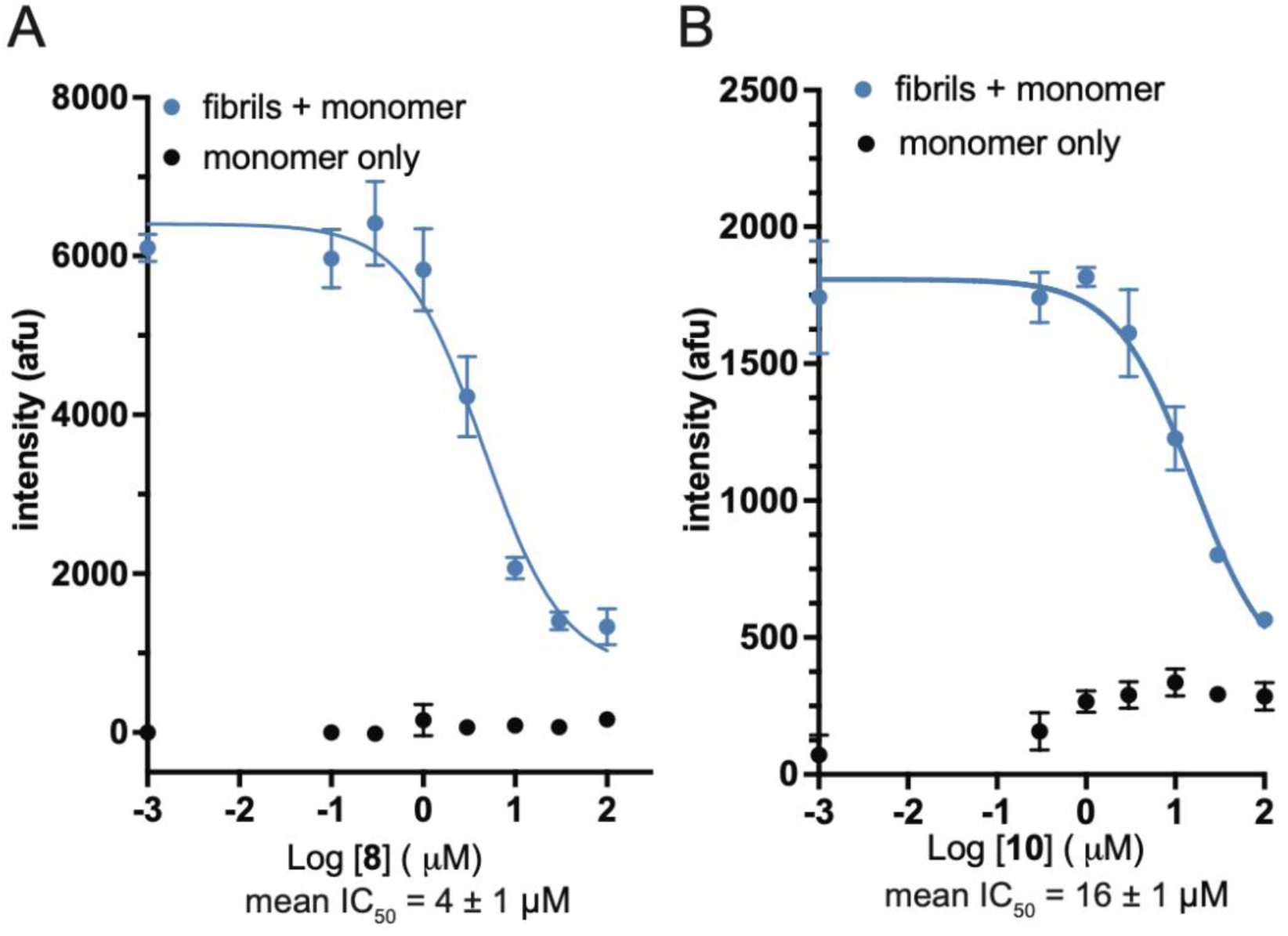
Fibril growth inhibition by the lead compounds is also observed for LBD-amplified fibrils. The dose-response curves of **8** (Figure 4A) and **10** (Figure 4B) in FlAsH fibril growth assays using fibrils amplified from LBD brain tissue. Graphs show mean (data points) plus std (error bars). Similar results were observed from two independent experiments.

### Docking studies suggest a potential mechanism for fibril growth inhibition

Small molecule compounds can potentially bind to fibrils or monomeric proteins in many different ways. When binding to the fibrils, the small molecule can bind to the highly ordered beta sheet region or, less likely, to the disordered N-terminal or C-terminal regions. To explore potential modes of interaction for inhibitor compounds, we utilized molecular docking software, Autodock 4.2^49^, to predict potential binding sites of the inhibitor compounds on LBD asyn fibrils (Figure 6, Supplementary Figure 5). We focused on LBD fibrils (PDB ID: 8a9l) as a docking substrate since this structure was determined for asyn fibrils extracted from postmortem human brain tissue. SSNMR studies indicated that the structure of amplified LBD fibrils (PDB ID: 8fpt) is highly similar to 8a9l, but were not able to resolve the structural conformation of the E46-V66 region^47,48^. A structural model for our in vitro asyn fibrils is currently being refined based on single particle cryo-EM and solid-state NMR data but is not yet available for docking studies. To maximize the effect of observing binding to fibril ends in parallel or perpendicular to the fibril axis, we limited our docking substrate to 5 fibril units, which is sufficient to also identify binding sites spanning multiple monomeric units along the fibril surface.

**Figure 6.**
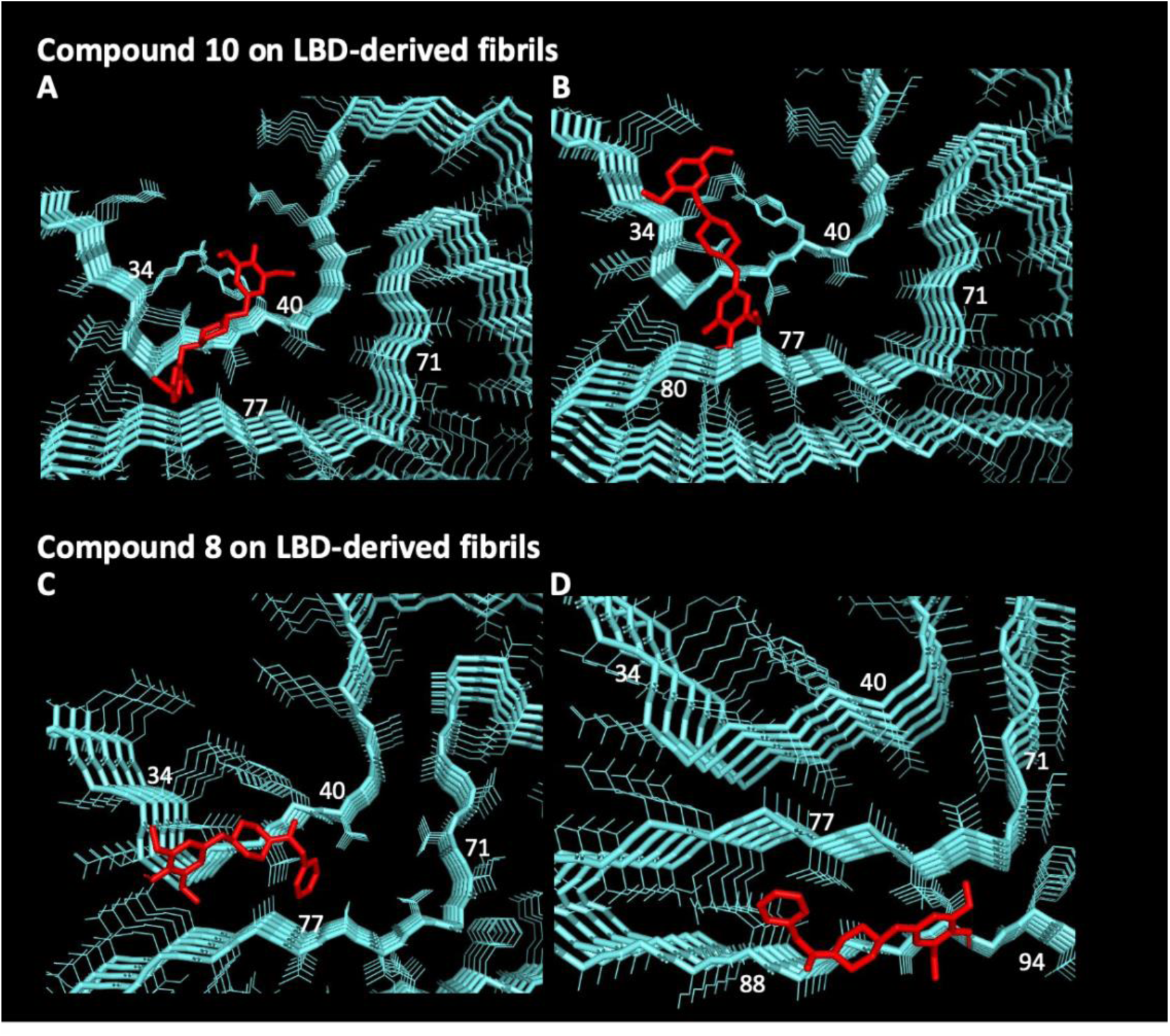
Binding sites predicted by molecular docking for compounds 10 and 8. In the examples shown in A-D, compound binding sites are on the surface of the elongating ends of fibrils, where they could inhibit binding of monomeric asyn required for elongation. Example docking sites of **10** (top 2 panels, A and B) and **8** (bottom 2 panels, C and D) on LBD asyn fibrils (8a9l).

Among binding sites with the lowest predicted binding energy for the two example compounds, we observed several poses on the surface of the elongating ends of the fibrils, which indicate a “capping” mechanism of inhibition where the compounds compete with the binding of monomeric asyn during elongation. Six of the nine docked sites for **8** and four of the nine sites for **10** are located on the surface of the elongating end of the fibril (Supplementary Figure 5). Interestingly for these binding sites, the compounds interact with one beta-strand and are oriented along the peptide backbone. The piperazine interacts with the backbone, while the aromatic groups on the left-hand and right-hand sides interact with the backbone and side chains. The interaction of the compounds with the elongating end may constrain the ability of additional monomers to align with the beta sheet conformation of the fibril seed, thus decreasing the rate of monomer association required for fibril growth. The alternative docked binding sites are along the fibril surface, parallel to the z axis, spanning 3 to 4 monomeric units (Supplementary Figure 5). Moreover, in some of these binding sites that are parallel to the z axis, the methoxy subgroups of the aromatic groups (left-hand or right-hand side) interact with fibril ends, which may also constrain the ability of additional monomers to attach to the elongating end.

## Discussion

This work optimized a fibril growth assay utilizing FlAsH fluorescence to screen and identify small molecule compounds that inhibit asyn fibril growth. The method enables the discovery of new compounds by measuring asyn fibril growth over short time periods (3 hours), using low concentrations of fibril seeds and monomer, where fibril growth is linear with respect to time^43^.

We obtained an IC_50_ value for EGCG in our FlAsH fluorescence assay that is similar to IC_50_ values previously obtained using ThioT to measure inhibition of asyn fibril growth^18,30^. Using our FlAsH fluorescence assay, we identified a family of dimethoxyphenyl piperazine.compounds capable of inhibiting fibril growth in-vitro. Fibril growth inhibition was further validated with both a ThioT fluorescence assay and an asyn conformation-specific immunoassay. The three assays produced similar results, but the FlAsH fluorescence assay can detect smaller changes in fibril growth over shorter periods of time and with greater precision. This is partly due to the absence of signal and noise from fibril seeds that do not contain C2-tags. The immunoassay and ThioT assay both detect initial fibril seeds as well as fibril growth, and hence, are less sensitive to small amounts of fibril growth that occurs in 3 hours. To compensate for these limitations, we increased the concentration of fibril seeds and monomer, and extended the incubation time (20 hours), in both the immunoassay and ThioT fluorescence experiments.

Our previous studies identified multiple locations where a single amino acid difference between monomeric asyn and fibril seeds can substantially alter the fibril growth rate^43,48^. Some amino acid residues within the beta-sheet region may be more important than others for determining the rate at which monomeric asyn associates with the elongating end of the fibril, since some amino acid changes had no effect on growth rates. These previous studies of fibril growth indicate that elongation requires specific conformational alignments as each monomeric unit associates with the fibril end. They also suggest that interaction of a small molecule compound with only two or three amino acid residues in either the fibril end or monomeric asyn has the potential to inhibit elongation.

Our docking studies indicate that dimethoxyphenyl piperazine compounds may inhibit fibril growth by binding to fibril ends at the elongating surface of fibrils and inhibiting the association of additional monomers. However, it is also possible that the compounds bind to monomeric asyn in ways that inhibit its association with fibrils or interfere with a conformational change required to associate with the fibril end. Other previously identified small molecules such as EGCG inhibit fibril growth by interacting with monomer and sequestering it through the formation of off-pathway oligomers to prevent association with elongating fibrils^23^. We observe an increase in FlAsH signal with incubation of monomeric asyn with EGCG, which is likely related to formation of oligomeric species. In contrast, FlAsH assays do not indicate the formation of oligomeric species when monomeric asyn is incubated with the dimethoxyphenyl piperazine compounds. Potential mechanisms based on interaction with either fibrils or momomer may be difficult to distinguish through further kinetic studies with varying concentrations of monomer or fibrils. For example, increasing monomer concentration may decrease the IC_50_ for compound inhibition by increasing the number of binding sites available on monomer or alternatively by promoting greater competition of monomer with compound at binding sites on fibril ends. Increasing fibril seed concentration may increase the number of binding sites available on fibril ends but may not change IC_50_ if it does not reduce free monomer concentration and also may not change the IC_50_ if compound is binding to monomer. Future studies using cryo-EM, NMR and radioligand binding assays with radiolabelled compounds of interest could provide additional information about binding sites and potential mechanisms. Further structural understanding of binding sites on fibrillar and monomeric asyn can guide the optimization of compound structure to improve potency.

This study’s limitation is that compounds were identified using in-vitro assays and have not been validated in cellular or in-vivo environments, where other mechanisms of fibril accumulation or clearance may be important. Studies of uptake and metabolism will also be required to optimize the potency of compound inhibition in cellular and in-vivo models.

To date, there have been several pre-clinical and clinical studies focusing on small molecules that prevent asyn fibril accumulation. EGCG, which has been shown to inhibit asyn fibril growth, failed to show a protective effect in participants with multiple systems atrophy, another synucleinopathy (phase 3, PROMESA trial). This could be due to the limited blood-brain-barrier penetration of EGCG^50,51^. Other small molecules currently in phase 2 trials include a pyrazino-pyrimidine NTP200-11 (UCB0599, NCT04658186) ^41,52^ and herbal ethanolic extract MT101-5 (Mthera Pharma Co, NCT06175767)^53^. There are ongoing efforts to study the tolerability of small molecules of a pyrazole moiety Anle138b(MODAG GmbH, phase 1 NCT04685265) that have been shown to decrease asyn aggregation in the presence of lipids^54,55^. Furthermore, there have been promising results from preclinical studies of small molecule compound SynuClean-D in vitro and in human neuroglioma cells ^9,10^ and with molecular tweezers (Clr01) in-vitro and in mouse models^56–59^.

With further optimization and validation, our assay may be suitable for high throughput screening of compound libraries to identify additional new leads for this PD therapeutic approach. Although the compounds identified here appear equally potent in assays with in vitro and LBD asyn fibrils, there may be advantages to utilizing asyn fibrils derived from postmortem LBD tissue in fibril growth assays as a primary screening approach for small molecule inhibitors. Our previous studies indicate that different asyn fibril conformers have distinct structural requirements for fibril growth, and identified several amino acid residues that are critical for LBD asyn fibril growth but not for in vitro fibrils. Future studies to understand how the compounds identified here and in other studies inhibit LBD asyn fibril growth may also provide further guidance for optimizing inhibition potency in efforts to develop effective mechanism-based therapeutics targeting asyn fibril accumulation in PD.

## Conclusion

Asyn fibril growth is a process that can be targeted to prevent the progressive accumulation of asyn fibrils in PD. Our assay provides a platform for identifying potent asyn fibril growth inhibitors. Ultimately, these small molecules could be further developed as disease-modifying therapies for PD.

## Methods and Materials

### Compounds

Compounds were purchased from Chembridge (San Diego, California USA) and Vitas-M Laboratory (Hong Kong). They were dissolved in DMSO at a concentration of 10mM and stored at −20°C.

### Preparation of recombinant WT-asyn, C2-WT-asyn and asyn fibrils

Untagged WT-asyn monomer, C2-tagged WT-asyn monomer, and asyn fibrils were prepared as described^43^. Briefly, asyn protein was expressed in E. Coli (BL21-CodonPlus (DE3)-RIL Competent Cells, Agilent Catalog: 230245) using the pRK172 plasmid expressing the asyn sequence. Bacteria were grown in TB broth containing 50 μg/ml ampicillin overnight with shaking. Asyn monomeric protein was extracted using osmotic shock followed by heat denaturation and ion-exchange chromatography. Purified asyn monomer was stored at −80°C until use.

WT-asyn and C2-WT-asyn fibrils were made by incubating 2 mg/ml of WT-asyn protein or C2-WT-asyn protein, respectively, in 20 mM Tris-HCl, pH 8.0 plus 100 mM NaCl buffer with shaking at 1000 rpm in an Eppendorf thermomixer set to 37°C. Fibrils were stored at 4°C until use. The concentration of asyn monomer and fibrils was determined using a micro-BCA assay performed according to the manufacturer’s instructions. Briefly, we utilized the micro-BCA assay (Thermo Fisher, Catalog number 23235) to estimate the amount of fibrils. A standard curve (200 µg/ml to 0.5 µg/ml) using manufacturer supplied diluted albumin (BSA) standard was used to determine concentration of asyn. The working reagent was made by mixing 25 parts of Micro BCA Reagent MA, 24 parts Reagent MB, and 1 part of Reagent MC (25:24:1, Reagent MA:MB:MC). In a microplate well, 150μL of each standard or unknown sample (three replicates) was added along with 150µl of the working reagent. The plate was incubated for 2 hours at 37 °C, cooled for 10 minutes and absorbance measured at 562nm on a plate reader. The amplified fibrils were centrifuged at 21,000g for 15 min at 4 °C to separate fibrils from monomer. The supernatant and total sample containing fibrils plus unincorporated monomer was also assessed. The measured decrease in asyn monomer concentration between total and supernatant was used to determine the concentration of fibrils.

### Preparation of LBD amplified fibrils

The Movement Disorders Brain Bank, Washington University, St. Louis, MO, provided well-characterized postmortem frozen brain tissue of de-identified participants. Written informed consent to perform a brain autopsy for research purposes was obtained from participants of the Movement Disorders Brain Bank. After death, the immediate next-of-kin were contacted and confirmed consent for brain removal and retention of brain tissue for research purposes. The participant who donated brain tissue used in these experiments participated in a clinical study approved by the Human Research Protection Office at Washington University prior to autopsy. All methods using human autopsy-derived material were carried out in accordance with relevant guidelines and regulations.

Fibril growth inhibition assays used amplified fibrils from case LBD1 as published previously^48^. Human postmortem brain tissue samples were sequentially extracted in buffers of increasing detergent strength using a Dounce Homogenizer to generate LBD-derived insoluble fraction seeds. The LBD seeds were combined with purified asyn monomer and subjected to 6 cycles of fragmentation followed by quiescent incubations to generate amplified fibrils.^48^

### Fibril growth assay utilizing FlAsH Fluorescence measurements

The fibril growth assay was performed similarly to previously published methods^43,48^. Briefly, fibril samples at 1μM concentration in fibril buffer were either sonicated for 5 min at amplitude 50 in a cup horn sonicator (Qsonica 600) for the recombinant WT- or C2- fibrils or 1 minute at amplitude 50 for the LBD-derived fibrils and mixed with 3μM of C2-asyn monomer and compound at a range of concentrations in 20 mM Tris-HCl pH 8.0, 100 mM NaCl buffer with 50 μL total volume. Compounds were dissolved in DMSO and the final concentration of DMSO was kept constant at 1.25%. The seeds, asyn monomer, and various concentrations of the compounds of interest were quiescently incubated for 3 hours at 37 °C in Corning Black 96-well plates (Fisher, catalog no. 07-200-762). After 3 hours, a 50 μL FlAsH assay mixture consisting of 7 mM tris(2-carboxyethyl)phosphine, 2 mM EDT, 2 mM EDTA, 50 nM FlAsH-EDT2 (Invitrogen TC-FlAsH II in-cell tetracysteine tag detection kit, catalog no. T34561), and 400 mM Tris-HCl, pH 8.0 was added and the total 100 μL mixture incubated for additional 1 hour at room temperature. FlAsH fluorescence was detected in a BioTek plate reader using a 485/20-nm excitation filter, a 528/20-nm emission filter, top 510-nm optical setting, and gain setting 100. Measurements were performed in triplicates. Dose-dependent curve fit a four-parameter logistic curve, Y=Bottom + (Top-Bottom)/(1+10^((LogIC_50_-X)*HillSlope)) to determine IC_50_. Mean IC_50_ was calculated from 2 to 3 experiments performed on different days. Maximum inhibition is calculated as (Intensity*_100 μM_*-Intensity*_0μM_*)/Intensity*_0μM_* x 100% where Intensity*_100μM_* and Intensity*_0μM_* is the intensity at 100μM of compound and intensity with no compound in the presence of fibrils and monomers.

### Measurement of quenching

Experiments to evaluate quenching were performed with 1 μM fibrils assembled from C2-WT-asyn plus 100 μM compound in the absence of monomer. Fibril seeds and compound were incubated for 30 minutes in 20 mM Tris-HCl pH 8.0, 100mM NaCl buffer (50 μL total volume), and then 50 μL FlAsH fluorescence assay mixture was added as above and incubated for an additional 1 hour at room temperature. FlAsH fluorescence was detected in a BioTek plate reader using a 485/20-nm excitation filter, a 528/20-nm emission filter, top 510-nm optical setting, and gain setting 100. Percent quenching is calculated as 100% - (Intensity_100 μM_/Intensity_0μM_).

### Measurement of native fluorescence

Experiments to evaluate native fluorescence were performed with 100 μM compound alone in the absence of fibril seeds and monomer. In a similar manner, compounds were incubated for 30 minutes in 20 mm Tris-HCl pH 8.0, 100mM NaCl buffer (50 μL total volume) and then 50 μL FlAsH fluorescence assay mixture was added as above and incubated for an additional 1 hour at room temperature. FlAsH fluorescence was detected in a BioTek plate reader using a 485/20-nm excitation filter, a 528/20-nm emission filter, top 510-nm optical setting, and gain setting 100. Percent native fluorescence is calculated as 100% x Intensity_100 μM_/intensity__DMSObuffer_, with Intensity_100 μM_ as the intensity of 100uM compound in the absence of monomer and fibrils and intensity__DMSObuffer_ is the background intensity of buffer.

### Sandwich ELISA

Sandwich ELISA was performed similarly to previously published methods^46^. Briefly, anti-alpha-synuclein antibody [MJFR-14-6-4-2] (Abcam ab209538) was used as capture antibody and diluted to 1 μg/mL in 34 mM sodium bicarbonate, 16 mM sodium carbonate solution with 3 mM sodium azide. The capture antibody solution was added to a 96-well plate (50 μL/well) and incubated overnight at 4°C. Plates were washed 5X with 150 μL 1XPBS + 0.05% Tween-20/well in between each step. Plates were blocked with 150 μL of 2% BSA (Sigma-Aldrich #A7906) in PBS at 37°C for 2 hours. Before standard and sample addition, plates were washed as previously stated. Samples were diluted at 1:10,000 in sample buffer containing no SDS to reach an SDS concentration of 0.06%. Standards and samples were added to the plate 50 μL/well in duplicate wells and incubated overnight at 4°C with slow rotation. The detection antibody (13G5B, in-house) solution was diluted to a final concentration of 1 μg/mL in PBS + 0.05% BSA + 0.05% Tween. The detection antibody solution was added 50μL/well and incubated at 37°C for 2 hours. Streptavidin poly-HRP80 (Fitzgerald # 65R-S118) was diluted 1:8000 in 1% BSA in 1XPBS+0.05% Tween-20. The streptavidin solution was added 50 μL/well and incubated for 90 minutes at room temperature in the dark. The plate was washed as described previously. Fifty μL/well of 3,3’,5,5’-Tetramethylbenzidine Liquid Substrate, Super Slow (Sigma # T5569) was rocked on the rotator until the first reading. Readings were done on Synergy 2 (BioTek) at 650 nM at 9, 12, and 15 minutes. The time point with the highest ratio between the top concentration and 0 concentration (background) in the standard curve was chosen for data analysis. Dose-dependent curve fit a four-parameter logistic curve, Y=Bottom + (Top-Bottom)/(1+10^((LogIC_50_-X)*HillSlope)) to determine IC_50_. Mean IC_50_ calculated from 2 experiments.

### ThioT fluorescence assay for fibril growth

The ThioT fibril growth assay was performed similarly to that of the FlAsH fluorescence assay with a few modifications. Fibril samples (3 μM concentration) were sonicated for 5 min at amplitude 50 in a cup horn sonicator (Qsonica 700) and mixed with 6μM of C2-asyn monomer and compound at 30 μM and 100 μM in 20 mm Tris-HCl pH 8.0, 100 mm NaCl buffer with 100 μL total volume. Compounds were dissolved in DMSO and the final concentration of DMSO was kept constant at 1.25%. The seeds, asyn monomer, and concentrations of the compounds of interest were incubated for 20 hours at 37°C in Eppendorf Thermomixer R at 1000rpm in ultracentrifuge tubes. After 20 hours, the assay mixture volume (100 μL) was increased to 300 μL with fibril buffer (20 mM Tris-HCl pH 8.0, 100 mM NaCl buffer) and centrifuged for 100,000×g for 20 min at 4℃. The supernatant (270 μL) was removed and replaced with 270 μL fibril buffer. 300 μL total volume was again centrifuged at 100,000×g for 20 min at 4℃. Again, the supernatant (270 μL) was removed, the seeds were recovered and mixed with 50 μL fibril buffer with 0.1% Triton. The 80μL sample was placed into a Corning Black 96-well plates (Fisher, catalog no. 07-200-762) and incubated with 18 μM ThioT in Fibril Buffer. A total volume of 160 μl/well was incubated at room temperature for 24 hours at 200 rpm in a Benchmark Scientific Incumixer. After 24 hours, ThioT fluorescence was measured in a BioTek plate reader using a 440/30-nm excitation filter, a 485/20-nm emission filter, and top 50% optical setting. % Fibril Growth without compound is calculated by (Intensity Measurement – Intensity_fibrils_only)/(Intensity_nocompound - Intensity_fibril_only), where Instensity_fibril_only is the average intensity of fibrils in the absence of monomer and compounds, Intensity_ nocompound is the average intensity of fibrils incubated with monomers in the absence of compounds.

### Molecular Docking

Compound structures were downloaded as 3D SDF files (Chembridge, Vitas M Laboratory). Molecular blind docking studies were performed via Autodock 4.2^49^ (Scripps) and viewed with Pymol (pymol.org). The structure of LBD-derived asyn fibrils determined by single particle cryo-EM (PDB ID: 8a9l)^47^ were obtained from RCSB protein data bank (https://www.rcsb.org/) as a target protein for blind docking. We decreased the LBD-derived asyn structure 8a9l to 5 fibril units, kept only the beta-sheet regions aa 31-100, and applied the docking structures. Water molecules and nonpolar hydrogens were removed from all compounds and protein structures. A grid box with a dimension of 126 x 126 x 126 Å^3^ was applied to all asyn structures, and center x= 86.868, y= 95.080, and z=97.452 were applied for 8a9l. Energy range was set to 4 and exhaustiveness was set to 8 (default values).

## Acknowledgements

We thank Rebecca L. Miller, Jennifer Y. O’Shea, and Aditi Bagade for helpful discussions. Support for this work was provided by: research grants from the Michael J. Fox Foundation and the American Parkinson Disease Association; NIH grants NS110436, NS097799, NS075321 from the National Institute of Neurological Disorders and Stroke and the National Institute on Aging; an American Academy of Neurology Clinical Research Training Scholarship in Parkinson’s Disease, funded by the Parkinson’s Foundation and the American Brain Foundation.

## Author Contributions

Conceptualization and Study Design: PTK, NJC, JMB, HH; Method development: HH, DD, SJW, PTK; Acquisition of data: HH; Analysis and interpretation of data: HH, DD, PK; Drafting of the manuscript: HH, DD, PK; Critical review and revision of the manuscript: HH, DD, PTK, NJC, SJW

## Data Availability Statement

The authors declare that the data supporting the findings of this study are available within the paper and its Supplementary Information files. Raw data files are available from the corresponding author upon reasonable request.

