## Supplementary Material for "Identification of Small Molecule Dimethyoxyphenyl Piperazine Inhibitors of Alpha-Synuclein Fibril Growth"

### Supplementary Information

**A**

| 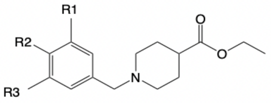 | R1 = | R2 = | R3 = | IC50 (µM) (mean ± std) | Max inhibition (%) (mean ± std) |
|---|---|---|---|---|---|
| 1 | OCH3 | OH | OCH3 | 18 ± 3 | 79 ± 4 |
| 2 | H | OCH3 | H | no inhibition | 16 ± 6 |
| 3 | H | OH | H | >100 | 30 ± 6 |
| 4 | OCH3 | H | OCH3 | no inhibition | 12 ± 2 |

**B**

| 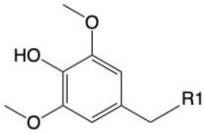 | Structure with R1 (shaded in grey) | IC50 (µM) (mean ± std) | Max inhibition (%) (mean ± std) |
|---|---|---|---|
| **5** | 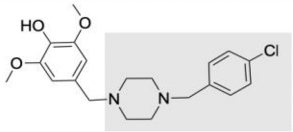 | 15 ± 2 | 99 ± 7 |
| **6** | 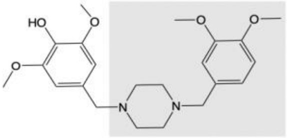 | >100 | 28 ± 15 |
| **7** | 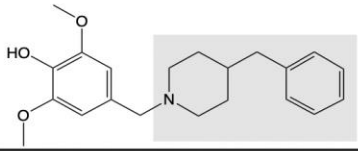 | no inhibition | 10 ± 4 |

Supplementary Figure 1

Supplementary Figure 1. Summary of SAR studies. A) Results of initial SAR studies (SAR1) show minimal asyn fibril growth inhibition with substitutions in the benzyl group (left-side) of **1**. $IC_{50}$ is mean ± std. Max inhibition is calculated as $(Intensity_{100\ \mu M}-Intensity_{0\mu M})/intensity_{0\mu M}$ x 100%. N =2 independent experiments. B) Results of SAR2 studies with substitutions to the right-side of **1**. $IC_{50}$ is mean ± std from 2 different independent experiments. Max inhibition is calculated as $(Intensity_{100\ \mu M}-Intensity_{0\mu M})/intensity_{0\mu M}$ x 100%.

| 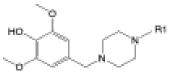 | R1 = | IC50 ($\mu$M) (mean ± std) | Max inhibition (%) (mean ± std) |
|---|---|---|---|
| **14** | 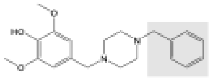 | 18 ± 1 | 95 ± 8 |
| **15** | 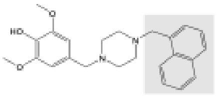 | 29 ± 2 | 71 ± 11 |
| **16** | 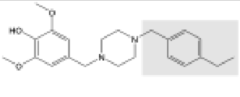 | >100 | 58 ± 15 |
| **17** | 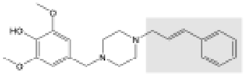 | >100 | 48 ± 20 |
| **18** | 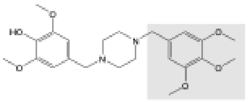 | >100 | 46 ± 12 |
| **19** | 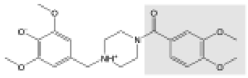 | >100 | 46 ± 2 |
| **20** | 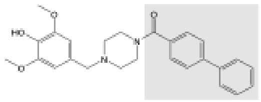 | >100 | 44 ± 20 |
| **21** | 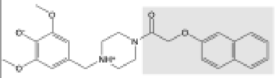 | >100 | 42 ± 6 |
| **22** | 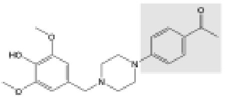 | >100 | 39 ± 4 |
| **23** | 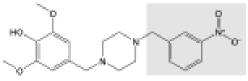 | >100 | 37 ± 5 |
| **24** | 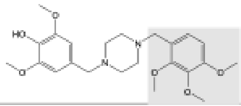 | >100 | 30 ± 5 |
| **25** | 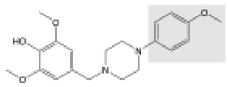 | No inhibition | 29 ± 2 |
| **26** | 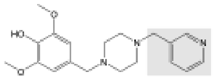 | No inhibition | 27 ± 2 |
| **27** | 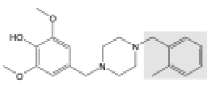 | No inhibition | 25 ± 4 |
| **28** | 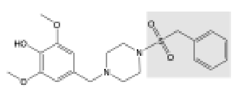 | No inhibition | 23 ± 14 |
| **29** | 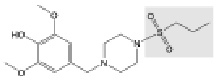 | No inhibition | 19 ± 6 |
| **30** | 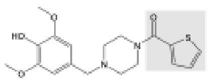 | No inhibition | 15 ± 4 |

Supplementary Figure 2. Results of SAR3 studies. $IC_{50}$ is mean ± std. Max inhibition is calculated as $(Intensity_{100\ \mu M} - Intensity_{0 \mu M})/intensity_{0 \mu M} \times 100\%$. N =2 independent experiments.

Supplementary Figure 2

### Supplementary Table 1

| Compound | % Fluorescence = 100% x $\frac{\text{Intensity\_100uM}}{\text{Intensity\_}DMSObuffer}$ (mean | ± | std) |
|---|---|---|---|
| **2** | 89.0% | ± | 0.2% |
| **3** | 90.0% | ± | 1.6% |
| **4** | 97.5% | ± | 2.1% |
| **5** | 99.2% | ± | 1.4% |
| **6** | 97.6% | ± | 1.7% |
| **7** | 99.4% | ± | 1.1% |
| **8** | 91.1% | ± | 0.0% |
| **9** | 94.2% | ± | 0.4% |
| **10** | 102.7% | ± | 0.2% |
| **11** | 100.3% | ± | 0.4% |
| **12** | 99.4% | ± | 0.9% |
| **13** | 95.8% | ± | 0.5% |
| **14** | 99.1% | ± | 0.3% |
| **15** | 108.1% | ± | 0.3% |
| **16** | 106.5% | ± | 1.0% |
| **17** | 94.1% | ± | 1.7% |
| **18** | 100.5% | ± | 0.7% |
| **19** | 102.2% | ± | 0.5% |
| **20** | 91.5% | ± | 1.4% |
| **21** | 97.0% | ± | 1.5% |
| **22** | 98.5% | ± | 0.0% |
| **23** | 91.5% | ± | 0.9% |
| **24** | 100.4% | ± | 0.7% |
| **25** | 101.7% | ± | 1.3% |
| **26** | 92.9% | ± | 0.7% |
| **27** | 103.6% | ± | 2.6% |
| **28** | 94.2% | ± | 0.1% |
| **29** | 97.4% | ± | 0.1% |
| **30** | 110.7% | ± | 6.0% |

Supplementary Table 1 . Native fluorescence measurements for the compounds. Calculated percent native fluorescence for compounds (relative to buffer) in the absence of preformed fibrils. Percent native fluorescence is calculated as 100% x Intensity$_{100\ \mu M}$/intensity$_{_DMSObuffer}$

Supplementary Table 2

| Compound | % Quenching = $100\% - \frac{Intensity\_100uM}{Intensity\_0uM} \times 100\%$ (mean | ± | std) |
|---|---|---|---|
| 2 | 9.2% | ± | 9.9% |
| 3 | 12.3% | ± | 0.9% |
| 4 | 9.8% | ± | 3.2% |
| 5 | 8.7% | ± | 3.0% |
| 6 | 25.8% | ± | 0.5% |
| 7 | 8.3% | ± | 1.8% |
| 8 | 7.0% | ± | 7.8% |
| 9 | 3.6% | ± | 7.3% |
| 10 | -3.2% | ± | 1.9% |
| 11 | 0.8% | ± | 2.7% |
| 12 | 11.3% | ± | 7.6% |
| 13 | 14.6% | ± | 3.5% |
| 14 | 5.1% | ± | 5.6% |
| 15 | 28.1% | ± | 0.8% |
| 16 | 9.2% | ± | 2.0% |
| 17 | 5.4% | ± | 0.9% |
| 18 | 8.7% | ± | 6.0% |
| 19 | 10.3% | ± | 2.7% |
| 20 | 11.1% | ± | 9.5% |
| 21 | 6.7% | ± | 5.1% |
| 22 | 1.5% | ± | 3.1% |
| 23 | 24.2% | ± | 7.2% |
| 24 | 2.3% | ± | 1.5% |
| 25 | 10.0% | ± | 2.6% |
| 26 | 7.2% | ± | 1.7% |
| 27 | 6.2% | ± | 1.5% |
| 28 | 17.2% | ± | 2.9% |
| 29 | 8.6% | ± | 6.6% |
| 30 | 1.4% | ± | 2.9% |

Supplementary Table 2. Fluorescence quenching measurements for the compounds. Calculated percent quenching for compounds in the presence of preformed C2-asyn fibrils and FlAsH dye. Percent quenching is calculated as 100%- ($Intensity_{100\ \mu M}/Intensity_{0\mu M}$)x100%.

### Supplementary Figure 3

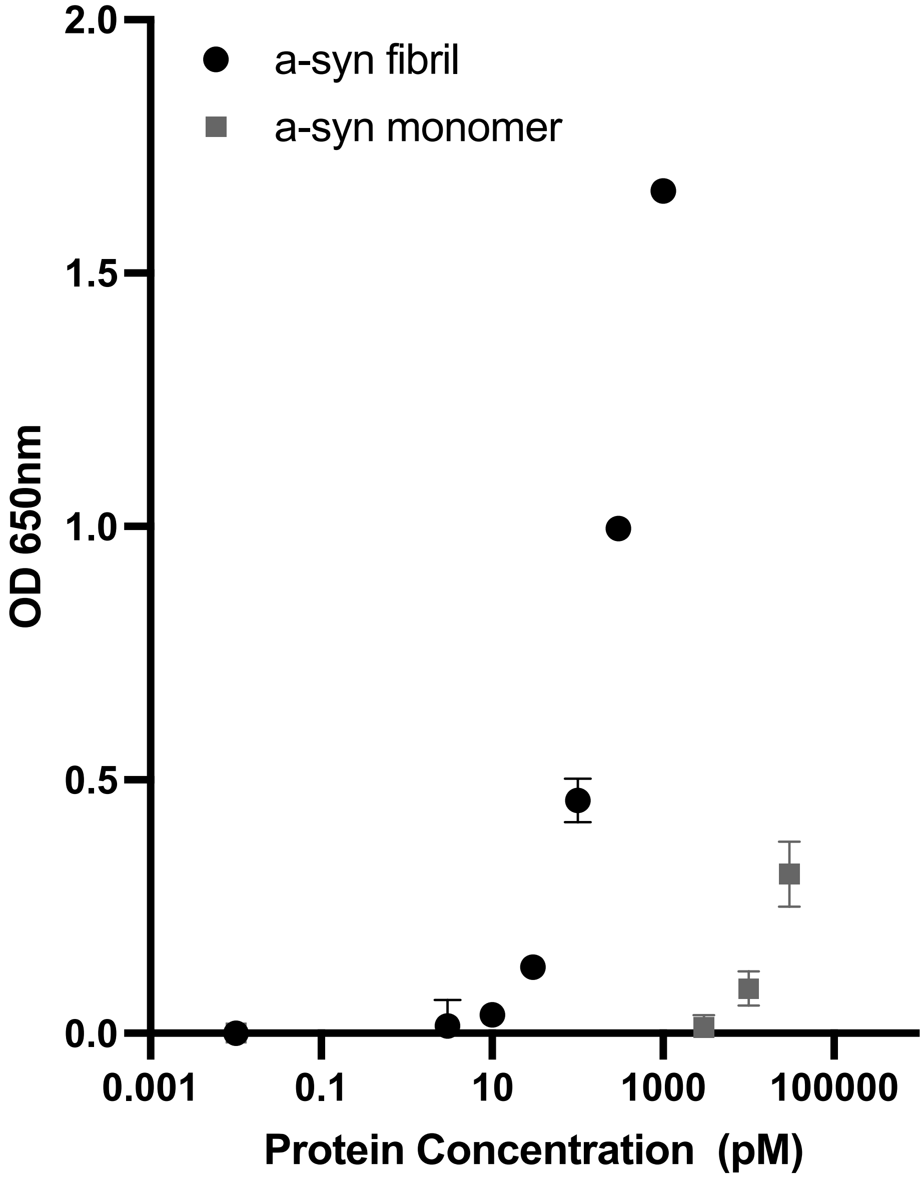


Supplementary Figure 3. Relative sensitivity of the immunoassay for fibrils and monomers. $IC_{50}$ is mean ± std. This assay is approximately 300-fold more sensitive for detecting fibrils than monomers.

### Supplementary Figure 4

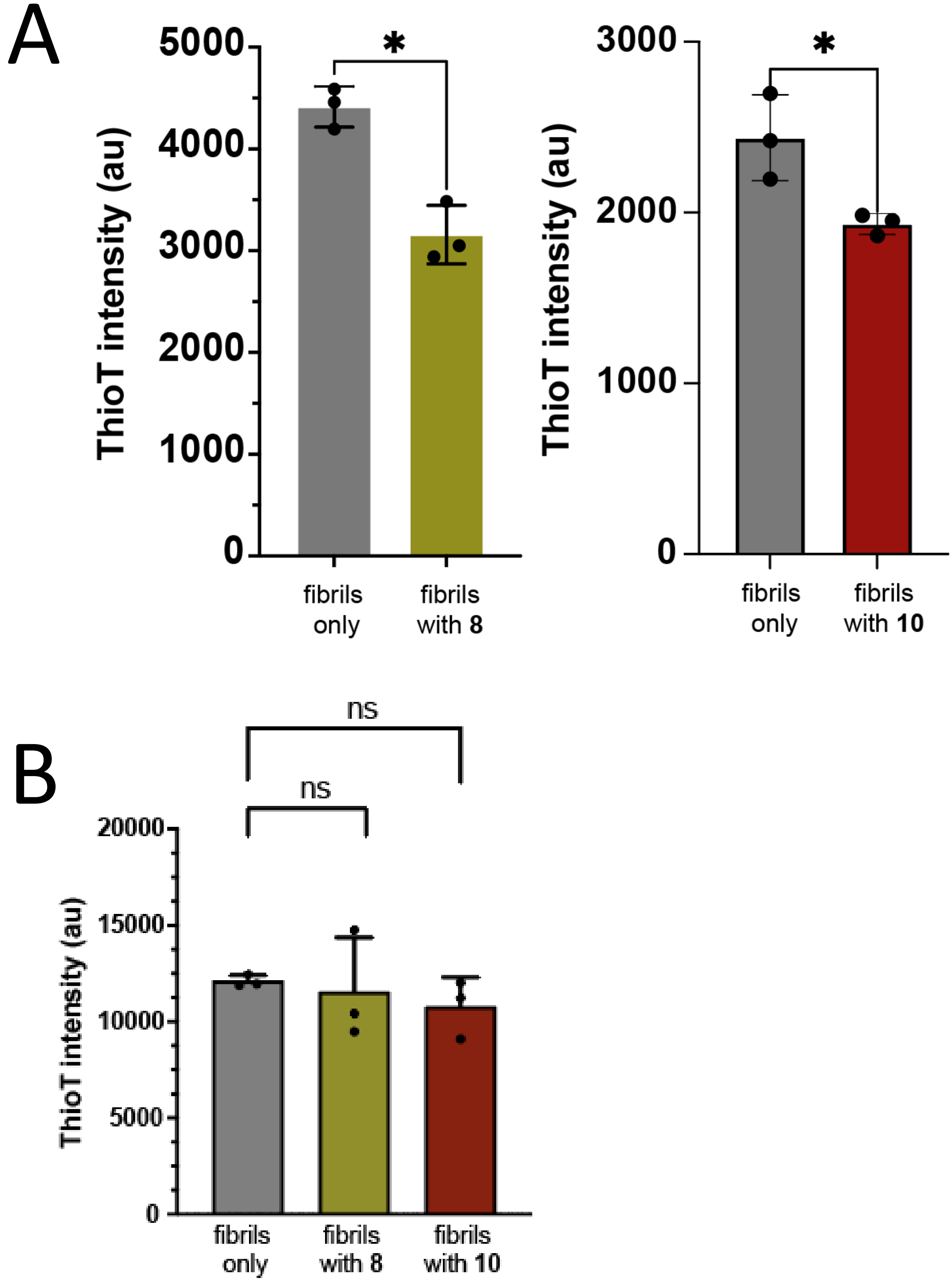


Supplementary Figure 4. A) Fluorescence intensity of ThioT plus fibrils without compounds (fibrils only) and ThioT plus fibrils in the presence of 100 μM of **8** and **10**, incubated in the absence of monomer. Reduced ThioT fluorescence signal is observed in the presence of **8 and 10**, (t-test, $p<0.05$). B) After serial wash and centrifugation to remove compounds **8** and **10**, there is no significant difference in ThioT fluorescence between the test conditions (t-test, $p<0.05$). Measurements were performed after incubation with DMSO buffer or 100 μM compounds. Data points represent mean ± std of triplicates. N= 3 three independent experiments.

A

Compound 8 Docked to LBD-derived fibrils (8a9l)

| Mode | Affinity (kcal/mol) | Image | Orientation to Fibril? (Parallel/Perpendicular) |
|---|---|---|---|
| 1 | -6.8 | 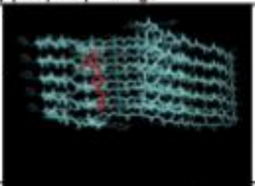 | Perpendicular |
| 2 | -6.7 | 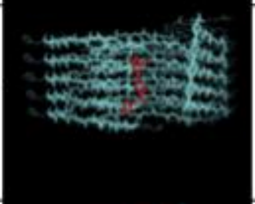 | Perpendicular |
| 3 | -5.9 | 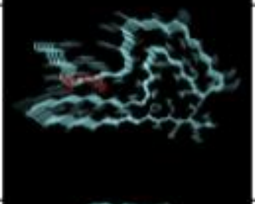 | Parallel |
| 4 | -5.8 | 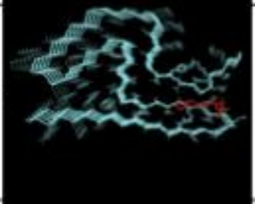 | Parallel |
| 5 | -5.6 | 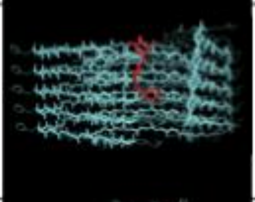 | Perpendicular |
| 6 | -5.5 | 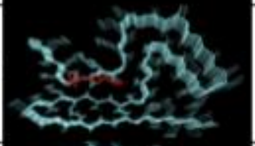 | Parallel |
| 7 | -5.2 | 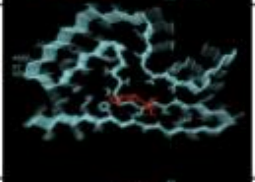 | Parallel |
| 8 | -5.2 | 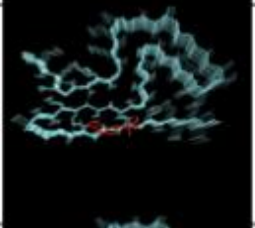 | Parallel |
| 9 | -5.2 | 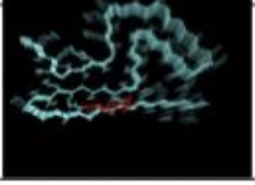 | Parallel |

B

Compound 10 Docked to LBD-derived fibrils (8a9l)

| Mode | Affinity (kcal/mol) | Image | Orientation to Fibril? (Parallel/Perpendicular) |
|---|---|---|---|
| 1 | -5.5 | 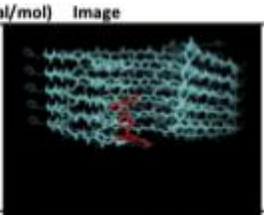 | Perpendicular |
| 2 | -5.3 | 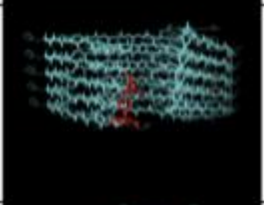 | Perpendicular |
| 3 | -5.2 | 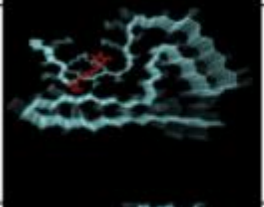 | Parallel |
| 4 | -5.0 | 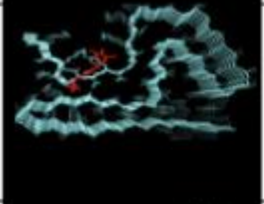 | Parallel |
| 5 | -5.0 | 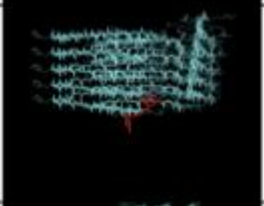 | Perpendicular |
| 6 | -4.7 | 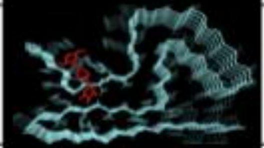 | Parallel |
| 7 | -4.7 | 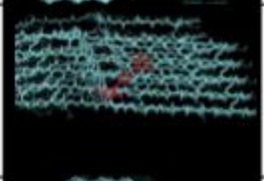 | Perpendicular |
| 8 | -4.7 | 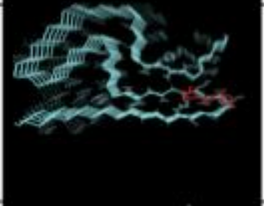 | Parallel |
| 9 | -4.7 | 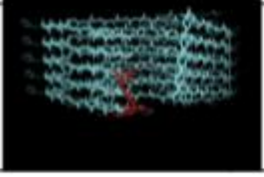 | Perpendicular |

Supplementary Figure 5

Supplementary Figure 5. Molecular docking results for **8** (left, A) and **10** (right, B) on LBD-derived fibrils (8a9l) with calculated free energy affinity for each predicted binding site.
